# Constrained standardization of count data from massive parallel sequencing

**DOI:** 10.1101/2021.03.04.433870

**Authors:** Joris Vanhoutven, Bart Cuypers, Pieter Meysman, Jef Hooyberghs, Kris Laukens, Dirk Valkenborg

**Affiliations:** Flemish Institute for Technological Research (VITO), Boeretang 200, B-2400 Mol, Belgium; Universiteit Hasselt, Data Science Institute (DSI), Interuniversity Institute for Biostatistics and Statistical Bioinformatics (I-BioStat), Agoralaan, Diepenbeek, BE 3590; Universiteit Antwerpen, Centre for Proteomics, Groenenborgerlaan 171, Antwerpen, BE 2020; Universiteit Antwerpen, Biomedical Informatics Network Antwerp (Biomina), Middelheimlaan 1, Antwerpen, BE 2020; Molecular Parasitology Unit, Institute of Tropical Medicine, Nationalestraat 155, Antwerpen, BE 2020; Universiteit Antwerpen, Adrem Data Lab, Department of Computer Sciences, Middelheimlaan 1, Antwerpen, BE 2020; Universiteit Hasselt, Data Science Institute (DSI), Theoretical Physics, Agoralaan, Diepenbeek, BE 3590

**Keywords:** Normalization, RNA-seq, transcriptomics, proteomics, multi-omics

## Abstract

In high-throughput omics disciplines like transcriptomics, researchers face a need to assess the quality of an experiment prior to an in-depth statistical analysis. To efficiently analyze such voluminous collections of data, researchers need triage methods that are both quick and easy to use. Such a normalization method for relative quantitation, CONSTANd, was recently introduced for isobarically-labeled mass spectra in proteomics. It transforms the data matrix of abundances through an iterative, convergent process enforcing three constraints: (I) identical column sums; (II) each row sum is fixed (across matrices) and (III) identical to all other row sums. In this study, we investigate whether CONSTANd is suitable for count data from massively parallel sequencing, by qualitatively comparing its results to those of DESeq2. Further, we propose an adjustment of the method so that it may be applied to identically balanced but differently sized experiments for joint analysis. We find that CONSTANd can process large data sets with about 2 million count records in less than a second whilst removing unwanted systematic bias and thus quickly uncovering the underlying biological structure when combined with a PCA plot or hierarchical clustering. Moreover, it allows joint analysis of data sets obtained from different batches, with different protocols and from different labs but without exploiting information from the experimental setup other than the delineation of samples into identically processed sets (IPSs). CONSTANd’s simplicity and applicability to proteomics as well as transcriptomics data make it an interesting candidate for integration in multi-omics workflows.

## Introduction

Any omics experimentation is prone to systematic effects (bias) introduced by latent or known factors, such as protocol and instrument variation. When the variation introduced by such a factor is of the same (or even higher) order of magnitude as the biological variation, then this effect must be accounted for. Otherwise, the biological effect under scrutiny can be obscured and wrong conclusions can be derived from the data. Normalization procedures or statistical analysis methods can be used to remove this bias.

RNA-Seq is a high-throughput technology used in transcriptomics that allows the relative quantification of thousands of RNA molecules simultaneously. A standard RNA-Seq workflow typically consists of the following steps: (1) Extraction, purification and quality control of the total RNA from the organism and experimental conditions of interest. (2) Purification of the relevant pool of RNA, e.g. enrichment of poly-A containing mRNA, or depletion of ribosomal RNA. (3) cDNA generation: the relevant RNA is converted to (more stable) cDNA using reverse transcription. (4) Library preparation and PCR amplification: adapters are added to the RNA/cDNA molecules of interest. These adapters typically contain sequences for compatibility with the sequencing platform, as well as barcodes to allow multiplexing of samples during a single sequencing run. After these adapters are added, we call the pool of molecules a ‘library’. If needed, the size of this library can be amplified with PCR. (5) Sequencing: sample libraries are pooled together and the molecules are sequenced in a massive parallel fashion. Millions of sequencing reads are generated. (6) Reads are demultiplexed if necessary and then bioinformatically mapped to the reference genome and gene read counts are summed per sample. (7) This data is merged to a count table, where the rows represent the genes, the columns the sample, and the values the gene read counts. This count table needs normalisation at two main levels before differential expression can be appropriately calculated: the library size and the experimental bias.

The first source of systematic variation, library size, corresponds to the number of reads for each sample. This number typically varies across samples, because it is impossible to add exactly the same number of molecules from each library to the sequencing instrument. The simplest way to correct for this would be to divide by the number of mapped reads, however, this can result in skewed differential expression results. Indeed, highly expressed genes have proportionally more reads, and if these vary systematically in expression level between conditions, they will disproportionately affect the number of reads, rendering the correction invalid. Hence, RNA-Seq data analysis software uses more sophisticated methods to correct for this. The core principle is always that the majority of genes do not change expression between samples. In DESeq2[1], this is done by first calculating the geometric mean read count for each gene across all samples. Then the read counts for each gene in a sample are divided by the mean across all samples to create a ratio. The final correction factor for each sample is the median of the ratios calculated for each gene in the sample. An alternative method, edgeR[2], uses trimmed mean of M-values (TMM) as its default normalization. Another common normalisation technique (though mostly used in micro-array processing) is Quantile normalisation, where the distributions of gene expression levels are made identical in statistical properties, across samples.

The second source of systematic variation, experimental bias, is caused by variables and confounders linked to the experimental particularities and design. For example, large experiments could be split over different days, labs or handling personnel which could all introduce unwanted sources of variation. In addition, the samples could be derived from a population of individuals with different age categories, different genetic backgrounds, etc. Most RNA-Seq analysis software can, however, account for this. For example, DESEQ2 and edgeR fit a negative binomial generalized linear model, which allows correction for covariates by explicitly adding them as factors to be fitted in the final model. As these factors need to be fitted alongside the biological variation, they are only available at the end of the statistical analysis. Moreover, their effects must still be explicitly applied to the partly normalized quantification matrix in case the researcher wants to use the latter to, for instance, make visualizations.

The subject of this paper is CONSTANd, which was originally built for the proteomics use case, presented by Maes et al.[3] as a method for the normalization of isobarically-labeled mass spectra that are being multiplexed in a pooled sample. Its main rationale is to correct for all sample-wise and peptide-wise systematic effects as well as scaling all quantification values to the same order of magnitude. To do so, it relies on three constraints: in the quantification matrix, all column sums should be equal, all row sums should be fixed to some value (identical across multiplexes) and this row sum value should be the same for all rows. Even though these constraints seem mutually exclusive, it is possible to suffise all three through constrained standardization (CONSTANd), a variant of the Iterative Proportional Fitting Procedure (IPFP) described by Deming and Stephan[4]. The second rationale of this normalization is that the quantification values in the same ‘run’ are identically affected by the same systematic variation, unlike those from other runs. As an immediate consequence of the two rationales, one can choose the constraint constants (row and column sums) such that normalized measurements from different runs may be directly combined for further analysis. Although the balance between conditions should be identical across such runs (e.g. 4vs2 cannot be combined with 3vs3), a novel adaptation detailed in this article does allow them to have different sizes (e.g. combine 4vs4 with 3vs3). Note that all of this is possible because CONSTANd can correct for both for the sample size and the experimental bias. Our main conjecture is that the CONSTANd normalisation method can also be applied on data from other types of - omics fields that do not strictly require multiplexing (because of excellent technical reproducibility). This will require a generalization of the ‘run’ from proteomics to an ‘identically processed (sub)set’ (IPS) of samples, where we explicitly exclude the systematic (biological) effect/treatment of interest from the concept of ‘processing’. In other words, samples in an IPS have been subject to identical experimental protocols that introduce little or no bias between them, so that the (biological) effect of interest ought to be the main source of systematic variation between the samples. In this manuscript, we demonstrate that CONSTANd is indeed suitable for RNA-seq experiments by applying it to three data sets of varying size, organism, and complexity. We also benchmark its performance compared to DESeq2, since it is considered the ‘golden standard’ and other methods have been shown to attain similar performance[5]. The quality of the normalisation is evaluated with principal component analysis (PCA).

## Results

We compare PCA plots and hierarchical clustering dendrograms of raw count data, DESeq2-normalized data and CONSTANd-normalized data from three different RNA-seq experiments. Note: In order to properly visualize data normalized for both library size and experimental biases by DESeq2, one first needs to explicitly remove the experimental bias by calling the ‘removeBatchEffect’ function, after applying the ‘DESeq’ function. The default DESeq2 PCA plot only accounts for library size normalisation, and would not be a fair comparison for DESeq2’s ability to correct for experimental biases.

### Leishmania (Cuypers et al.[6])

The *Leishmania* data set was generated in-house on a single instrument, in a single run. A total of 16 samples were prepared by two different library preparation methods, spliced leader sequencing (SL) and Illumina TruSeq (IL), each contributing an IPS of 8 samples. Each IPS contains 4 samples of each of two life stages (LOG and STAT) of the same *Leishmania* strain.

Figure 1 shows how both DESeq2 and CONSTANd succeed at removing the experimental bias that is present in the raw data due to the different library preparation methods, and uncover the biological differences between the two life stages.

**Figure 1:**
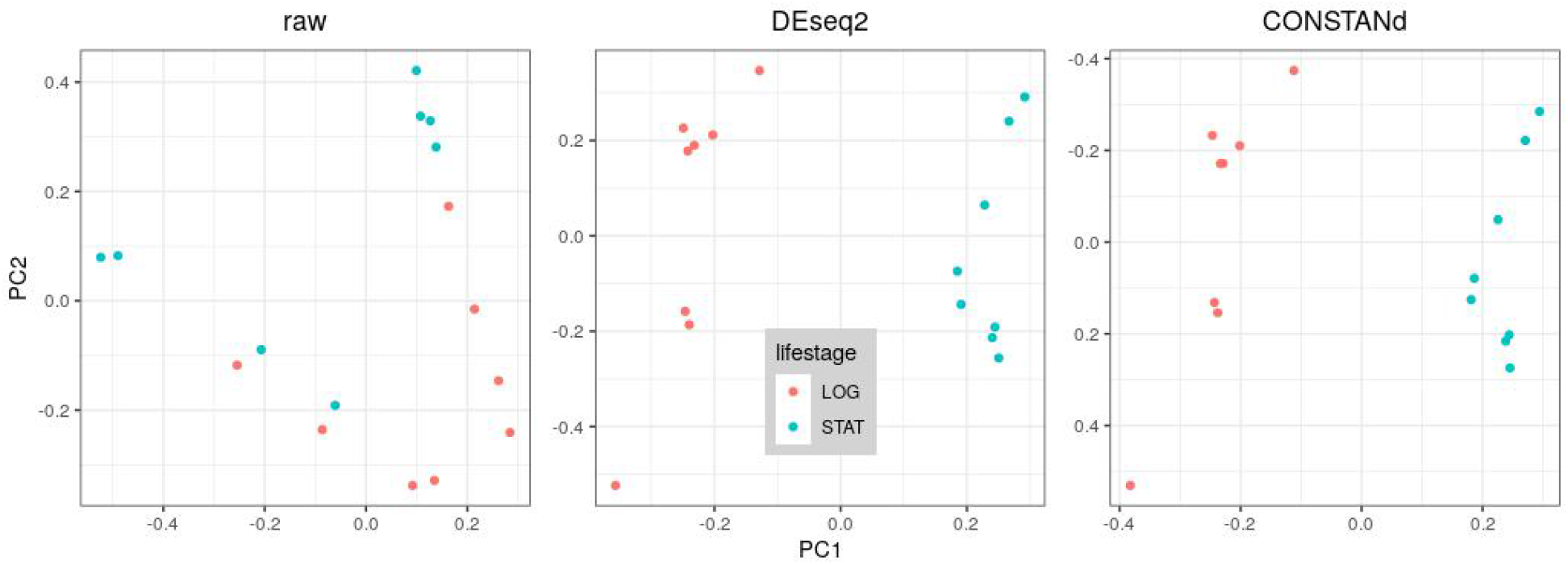
While the PCA does not group raw (non-normalized) data samples in the *Leishmania* data set (although they are separable according to library preparation method, see Fig. S5), both DESeq2 (after also correcting for experimental bias) and CONSTANd succeed at separating them according to their life stage on PC1. The PC2 axis in the CONSTANd plot was inverted to increase comparability.

### Mouse (Sarantopoulou et al.[7])

The Mouse data was taken from a publicly available data set generated on multiple runs, in a single site (lab). A total of 18 samples were prepared using three different library preparation methods (Pico, V4, TruSeq), corresponding to three separate runs, each contributing an IPS of 6 samples. Each IPS contains 3 samples (technical replicates from those in the other two IPSs) of each of two treatments (ILB and UNT).

Figure 2 shows how both DESeq2 and CONSTANd succeed at removing the experimental bias that is present in the raw data due to the different library preparation methods, and uncover the differences between the biological groups. This result was unexpected, because Sarantopoulou et al. performed the same analysis using DESeq2 and did not find grouping based on condition. However, in the Supplementary Information we argue that they have probably accidentally used non-normalized quantification values during some of their visualizations.

**Figure 2:**
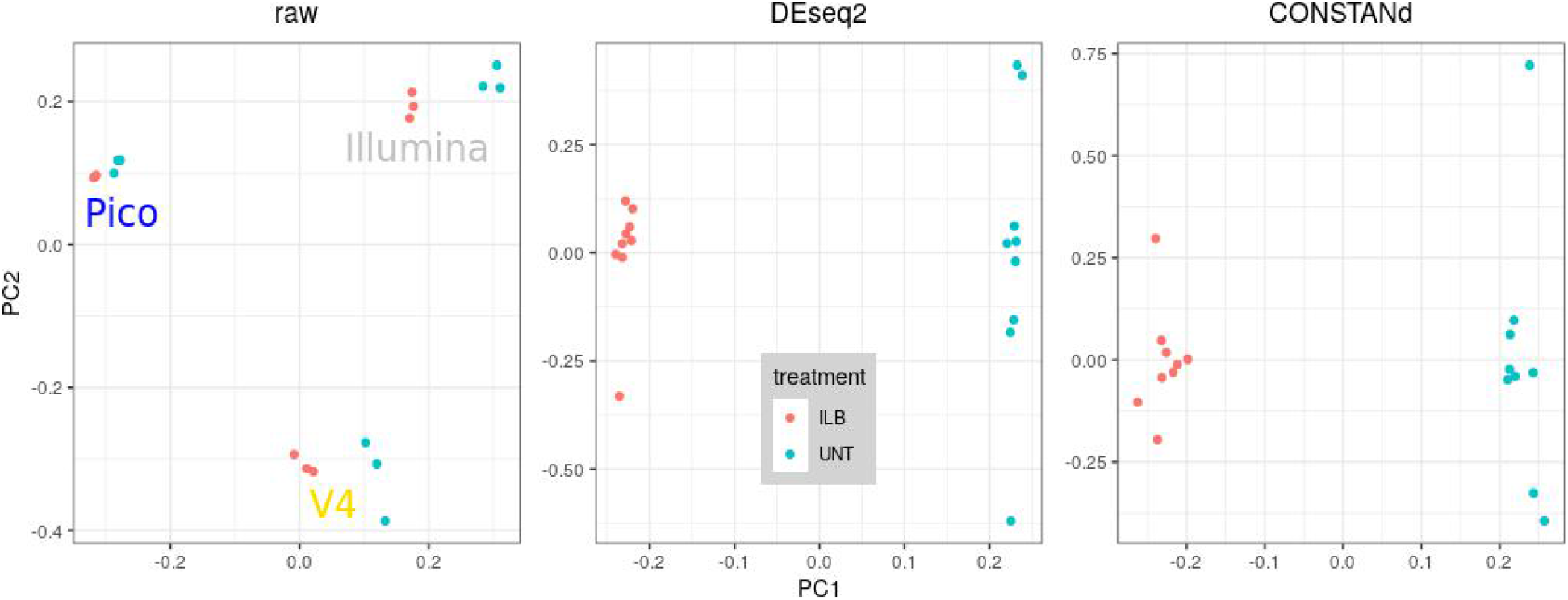
While the PCA groups raw (non-normalized) data samples according to library preparation method (see Figure S2), both DESeq2 (after also correcting for experimental bias) and CONSTANd succeed at separating the samples in the Mouse data set according to their biological condition on PC1.

### Human (Li et al.[8])

The Human data was taken from a publicly available data set generated by multiple runs, in multiple sites (L, R, V, W). A total of 50 samples were prepared, of which 32 in site W using two different library preparation methods (RiboDepletion, PolyA). Each lab contributes to an IPS of 6 samples (all RiboDep), except for lab W which gives rise to two IPSs (corresponding to RiboDep and PolyA): 16 samples each. Each IPS contains 3 samples (technical replicates from those in the other two IPSs) of each of two biological tissues A (pooled tumor) and B (brain tumor), except for the IPSs from site W which contain 4 samples of each A, B, C=¾A+¼B, D=¼A+¾B.

Figure 3 shows how both DESeq2 (runtime: about 120s) and CONSTANd (runtime: <1s) largely succeed at removing the site and library preparation bias that are present in the raw data (see also Fig.S6 and Fig.S7), and instead highlight the differences between the biological groups. Even though sample types C and D were present in only one site, this did not throw off either DESeq2 nor CONSTANd. All sample types A, D, C, B are spread out along the first principal component axis as one would expect based on their constituent fractions of pooled tumor and brain tumor (1:0, 0.75:0.25, 0.25:0.75, 0:1, respectively). In this case, the samples from site W were also split into two IPSs according to library preparation type for CONSTANd normalization. However, in the case where we normalize them together, that secondary experimental bias is still subordinate to the biological effect, as discussed in the Supplementary Information. The variability in samples from site W is noticeably larger than those from other sites, but Fig. S8 proves that this is not due to the presence of sample types C and D.

**Figure 3:**
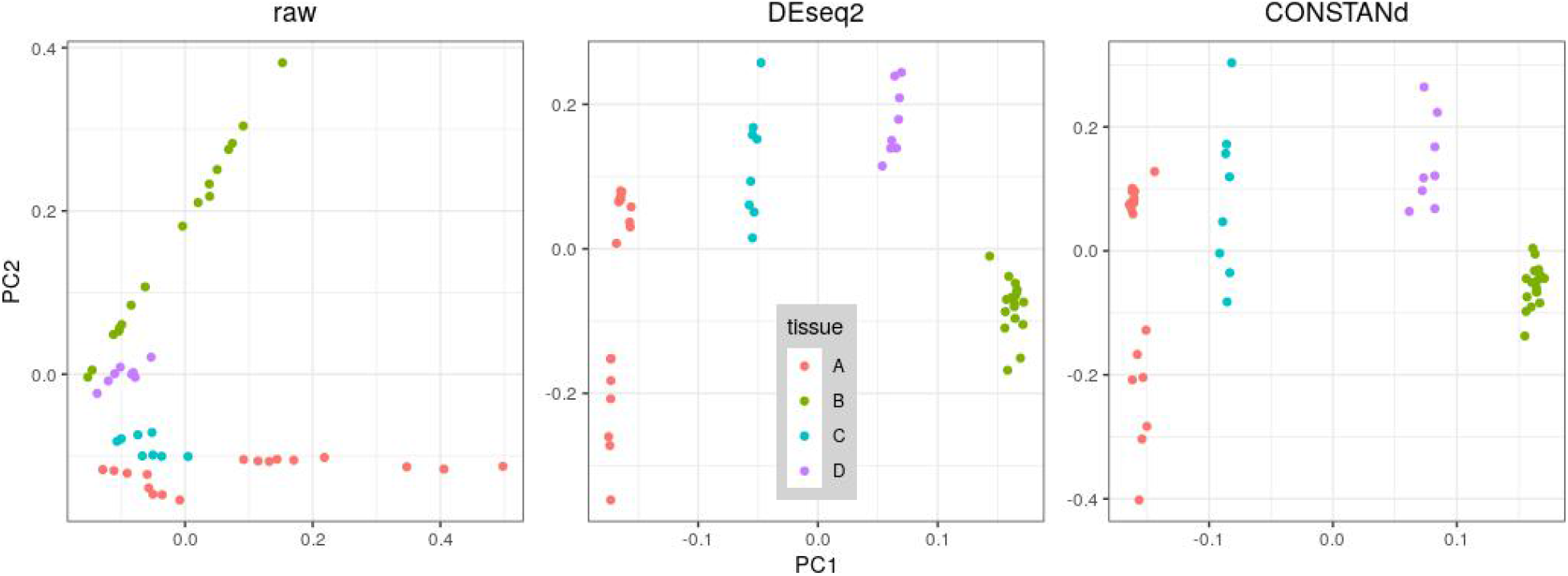
While the PCA only partially and only weakly groups raw (non-normalized) data samples according to all influential factors (see als Fig. S6 and Fig. S7), both DESeq2 (after also correcting for experimental bias) and CONSTANd succeed at separating the samples in the Human data set according to their biological condition on PC1.

## Discussion

In this manuscript, we illustrated that CONSTANd - an algorithm initially conceived to normalize label-based, multiplexed proteomics data - can also operate equally well on RNA-seq data. We illustrated this through analysis of three data sets of varying size, organism and complexity, and through comparison of the results with those of DESeq2, the current golden-standard software in the RNA-Seq field. All of our results clearly suggest that DESeq2 and CONSTANd are successful and comparable in their ability to properly normalize RNA-seq count data by removing all systematic bias while leaving effects of biological interest intact.

The results depicted in Fig. 3 and Fig. S1 of the Human data set indicate that it is indeed appropriate to delineate CONSTANd’s IPSs according to putatively influential latent factors in the data like site and library preparation method. In practice, these would be exactly those factors which one would use in the statistical model required by DESeq2, so this does not require any additional effort. In any omics experiment based on relative quantitation, there is always a level in the experimental set-up at which the samples can be put into IPSs. For labeled proteomic data, this level is defined by the pool of samples that are multiplexed. For other, potentially non-multiplexed types of data, this level is not clearly defined, but depends on the factors that introduce experimental bias in the data. One word of caution: even when technical reproducibility of instrumentation is excellent, a maintenance event or change in consumables may trigger a systematic effect. In such cases, still, the samples measured prior and after this event can be considered as two IPSs.

We deem CONSTANd is very easy to use for the following four reasons.

First, it performs all necessary data transformations in one procedure: the library size correction, the removal of experimental bias, and a magnitude-rescaling whose effect on the variance resembles that of vst. This stands in contrast with DESeq2, where each step requires a separate action by the user. Unifying these steps prohibits users from making mistakes, which is a non-negligible feature as we see demonstrated here by the putative mistake we have found (see Supporting Information) in the report by Sarantopoulou et al.[7], published in Nature Scientific Reports. We leave the assessment of any scientific implications up to the authors.

Second, CONSTANd’s methodology and implementation are extremely straightforward to understand: matrix raking as opposed to the statistical framework behind (generalized) linear models (GLMs). Even though the library size correction in DESeq2 is comparable in simplicity, any experimental biases (such as batch effects) need to be explicitly modeled in the GLM.

Second, Third, the normalization of different IPSs is done entirely independently. Therefore, in case one has an experiment spanning multiple instrument runs or even labs, one could immediately use CONSTANd after the data acquisition of the first part of the experiment as a QC step. Once all data has been collected and individually normalized, one only has to merge the output tables. This stands in contrast with model-based approaches like DESeq2, where one would have to re-specify the statistical model at each step and re-run the entire computation if one wants to do such preliminary analyses in addition to a final analysis. This is a step towards a faster analysis of massive parallel sequencing experiments.

Fourth, we have shown that CONSTANd is fast: its total runtime on all data of the Human data set was less than one second, which is more than two orders of magnitude shorter than the time it took DESeq2 to get to the point where a meaningful visualization of normalized data could be produced. This speed is a critical trait for any method that aims to quickly triage data sets according to quality and added value in a high-throughput field of study like transcriptomics and other - omics disciples alike.

Taking everything above into consideration, we believe that this normalization method would also be an excellent candidate for use in other - omics disciplines. CONSTANd’s putative applicability to these disciplines could enable the uniformization of the normalization layer in multi-omics studies.

## Methods

### CONSTANd: a method borrowed from different fields

CONSTANd is a data-driven normalization method for relative quantification experiments developed by Maes et al.[3]. Initially, CONSTANd was tailored towards the needs of multiplexed experiments in mass spectrometry-based proteomics. In this set-up, all proteins are digested in the wet lab and the resulting peptides are marked by an isobaric label whose type is unique for each sample. The samples are then multiplexed in a pool that is then measured and further analyzed. Three important requirements are derived from the nature of the scientific question (studying differential expression between proteomes) and the restrictions of wet lab procedures (requiring multiple instrument runs). Note that the manuscript by Maes et al.[3] mentions only two requirements because the third was included in the second.

First, to make quantification values comparable between different samples within a multiplex, we need to take into account that each is measured in proportion to the total size of that sample’s proteome. We are not interested in how many peptide or protein copies were in the test tube, but rather we would like to know whether there would be more or less of each type when comparing between equally-sized proteomes corresponding to different conditions of interest.

Second, in order to compare these values between different multiplexes, we need to also make sure they are not dependent on any processing steps that influence which parts of the proteome are measured and to what extent. Therefore, we need to also express each quantification value of a single peptide within a sample as a proportion of the total abundance of that peptide across all samples of the same multiplex, and make these total abundances uniform across multiplexes. Third, we do not want any protein to make disproportionate contributions to the proteome size for which we will correct (first requirement), nor to any visualizations (e.g. PCA plot) we will make. For instance, having a high quantification value is not equivalent to being abundant in-vivo since some peptides/proteins are more prone to getting measured than others. Therefore, we will rescale the magnitude of all observations by fixing all total abundances from the second requirement to the same value.

These three requirements are translated into constraints on the quantification matrix as follows:

1. For a particular sample: all intensities across the list of peptides should sum to *m/n* with *m* the number of peptides (rows) and *n* the number of multiplexed samples (columns).
2. For a particular peptide: all intensities across samples from the same multiplex (rows) must sum to the same fixed value across all multiplexes.
3. These row sums which are fixed across multiplexes should also be fixed to the same (arbitrary) value across all rows in each multiplex, which we choose to be one.

The careful, more mathematically inclined reader may notice that subjecting the quantification matrix to the first constraint prohibits it from satisfying the second and third, and vice-versa. Remarkably, though, there are several ways to circumvent this and solve such a constrained optimization problem, but CONSTANd employs this principle of ‘matrix raking’. In econometry this method is known as the iterative proportional fitting procedure (IPFP[4]) or also as the RAS procedure. In fact, RAS is a specific, very simple and computationally efficient implementation of the IPFP, first described by Stone et al.[9]. In fact, the pseudo-code of CONSTANd is depicted in Fig. 4 and comprises only a handful lines of code. The procedure iteratively rakes the original data matrix by each step re-scaling all values in each row or column so that their mean equals the ‘target’ value *1/n*, until the L1-norm of the difference between consecutive **K** matrices is small enough. The convergence is log-linear, so that typically after 10 to 20 iterations one obtains the final, normalized matrix **K**.

**Figure 4:**
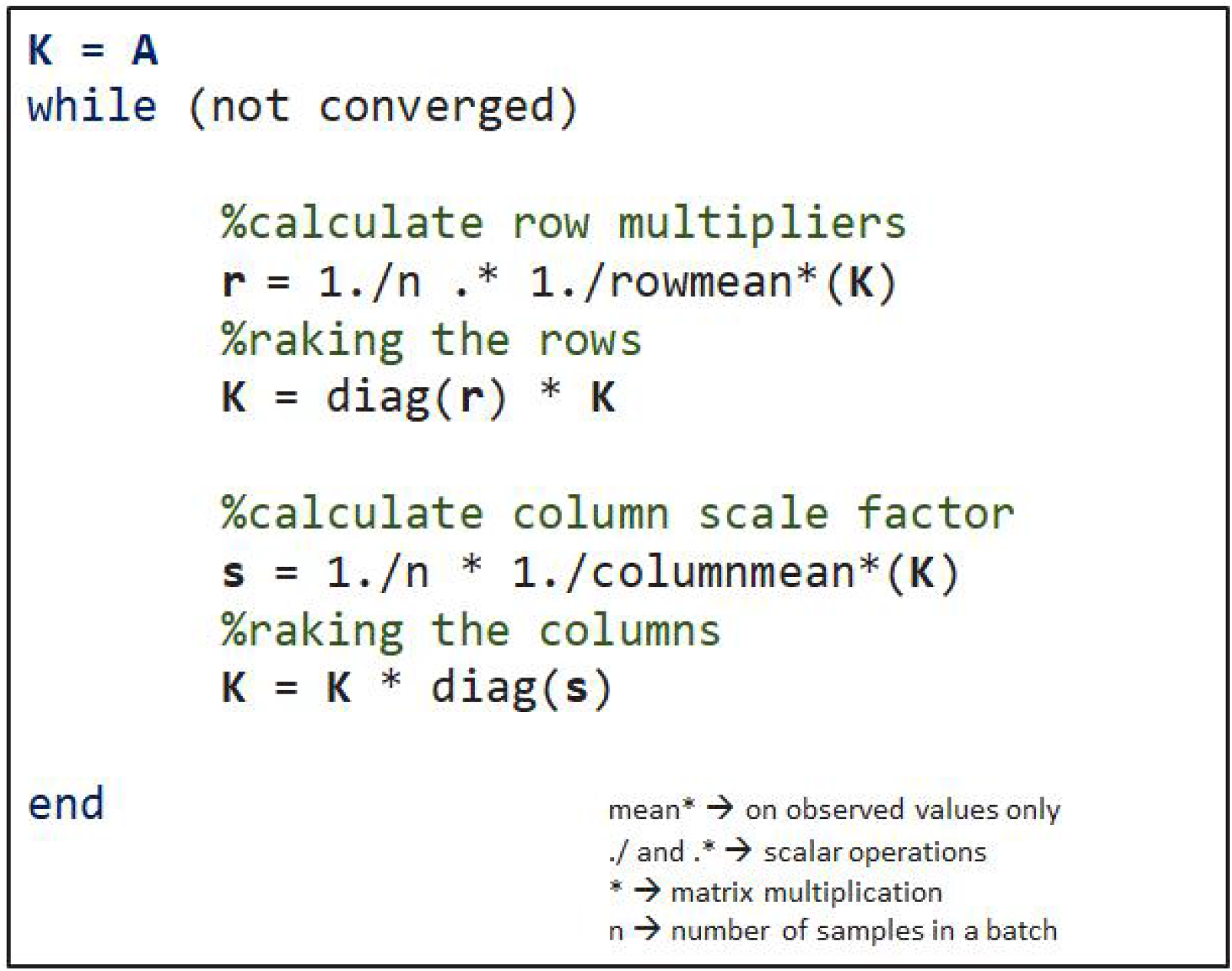
The CONSTANd variant of the RAS implementation of IPFP is very simple: it takes up only a handful lines of code. The mean here is defined as the ‘nanmean’, which ignores missing values. The authors believe a picture can say a thousand words even if, ironically, in this case it exists of nothing but a handful of words.

There are three assumptions the input data must adhere to. These apply to most data-driven normalization algorithms, but for completeness we mention them in a nutshell: (I) Most peptides/proteins are not differentially expressed; (II) The amount of overexpression in the data is roughly equal to the amount of underexpression; (III) The magnitude of any systematic bias is directly proportional to the intensity of the measurement. Further mathematical details are available in Maes et al.[3], and the up-to-date (see next section) online implementation of CONSTANd as well as detailed documentation are available at qcquan.net, where it also appears as part of the QCQuan proteomics workflow[10].

### CONSTANd: now applicable to differently-sized runs

This rationale of CONSTANd is sensible in case of labeled data and multiplexed samples as laboratories have only limited freedom to choose the size of the labeling-cassette. The disadvantage of the ‘original’ CONSTANd described above is that one cannot jointly analyze experiments that are balanced (as they ought to be) but are differently sized, e.g., 3vs3 samples in a run versus 5vs5 samples in a run. This is because the row and column means of the different experimental cassettes were constrained to *1/n* for different values of n (6 and 10, respectively). A straightforward, yet effective fix for this problem is to make the method invariant to the number of columns in a run by setting the row and column mean target value equal to 1, or in other words fixing *n*=1 in the constraints, regardless of experiment size. This allows one to first separately normalize and then jointly analyze multiple experiments of different sizes, as we demonstrate on the Human data set in this manuscript.

### Translation to a transcriptomics context

Transcriptomics data is generated in a completely different way then proteomics data. Despite, there are quite some similarities in the data that comes out of both technologies. Both are semi-quantitative - omic profiling technologies which in the end generate an - omics matrix in which the columns represent the samples, the rows the transcripts/proteins, and the intersection a quantification. In proteomics, these quantifications are mass-spectrometry based intensities, while in RNA-Seq these are read counts obtained by massive parallel sequencing. Importantly, in both disciplines a very common type of scientific question is that of differential expression between two or multiple types of -omic profiles, which requires no absolute but only relative quantification (with which CONSTANd is compatible). In quantitative proteomics, unless one wishes to use reference samples, multiplexing and labeling is an absolute necessity, because the technical variability at platform level (due to, e.g., slight variations in a peptide’s elution profile in the LC column, during LC-MS) between all sample measurements is enormous. In transcriptomics, however, RNA molecules are converted to small DNA fragments during library preparation. In contrast to peptides, these have very similar and reproducible properties, so that they do not meaningfully affect the experimental outcome. Therefore, even an entire series of RNA-seq sequencing runs may under the right conditions be treated in a similar fashion as the multiplexed proteomics samples from a single run. By ‘right conditions’ we mean that there exists no significant source of variability that may distinguish one sample from the other; they form an identically processed set (IPS). A counterexample: samples prepared using a different library preparation protocol, or the same library prep method at different moments, would *not* be in the same IPS. Other possible influential factors include ischemic time, centrifuge time, day effects, type of instrument, maintenance, etc. In general, the factors that delineate each IPS are exactly those that (might) constitute an additional source of (significant) variability. Although it is difficult to record all these factors, it is reasonable to treat samples that are measured together over a short period of time with the same protocol as belonging to the same IPS. In conclusion, we may apply CONSTANd separately to each count matrix corresponding to a specific IPS (which may span multiple runs), and afterwards re-combine them for joint analysis.

### Evaluation procedure

In order to evaluate whether the updated CONSTANd method is suitable for normalization of RNA-seq count data, a comparison with the state-of-the-art method DESeq2 is performed on three selected data sets. We also compare the result of the normalisation with the raw, unnormalized data. The measure of merit used for this comparison is based on the visual separation of the samples in PCA plots and/or hierarchical clustering dendrograms. The selected datasets are chosen because of the presence of three characteristics. First, the data sets are subject to an effect that is of scientific interest, like a treatment effect or biological condition. Second, there are biological replicates corresponding to this factor of interest that ought to group together. Third, there are one or multiple pronounced experimental biases due to differences in processing steps (e.g., different library preparation or instrument) that might jeopardize the grouping of these biological replicates. In a principal component analysis it is the idea to project the high-dimensional data set in an unsupervised manner onto a lower-dimensional space so that the information (variation) in the data becomes more pronounced. In an ideal world the PCA plot, which projects the data onto the two most information-rich dimensions, would reveal that biological replicates cluster together. The first principal component would separate the clustered biological replicates in relation to the effect of interest, whilst any experimental bias is removed from the data. Instead, without removal of such processing effects one risks grouping of the biological replicates according to the different processing factors they correspond to. A completely analogous rationale applies a fortiori to the interpretation of the hierarchical clustering dendrograms (with Euclidean distance metric), because they are a representation of information/variability from all principal components instead of just the first (though most important) two.

For all data sets in this manuscript, before normalization, we removed genes for which half or more of the count records were equal to zero. Any raw, unnormalized data were scaled by the R software’s ‘prcomp’ function when making the PCA plot. Similarly, any DESeq2-normalized data had their experimental bias explicitly removed using the ‘removeBatchEffect’ function from the ‘limma’ R package, and underwent a variance stabilizing transformation using DESeq2’s ‘vst’ function, but were not scaled by prcomp. Any CONSTANd-normalized data were not further modified before making plots, except for merging the count tables from multiple IPSs and zero-imputating any remaining missing values.

## Materials

### Leishmania data (Cuypers et al.)

A first data set was generated in-house and comprises 16 NGS samples as described in Cuypers et al.[6] that exhibit a severe experimental bias. A bias was artificially introduced when evaluating two alternative protocols for RNA library preparation. Eight *Leishmania* samples from two different growth stadia (LOG and STAT) were prepared with two library preparation methods and analyzed on the same NGS device. The library preparation methods spliced leader sequencing (SL) and Illumina TruSeq (IL) each delineate an IPSto which we apply CONSTANd separately, while the two life stages LOG and STAT are considered of biological interest. The same factors were specified in the model used by DESeq2. Any other remaining factors are ignored, because they would be confounded with a library preparation method.

### Mouse data (Sarantopoulou et al.)

In their comparative study of library preparation methods, Sarantopoulou et al.[7] performed RNA-Seq on liver samples from six twelve-week old male C57/B6J mice. Three were treated with IL-1β (ILB) and the other three with saline (UNT) by intraperitoneal injection. They then “*performed RNA-seq on those 6 samples using three different library preparation kits. […] (1) **Pico**: […] Takara Bio SMARTer: SMARTer Stranded Total RNA-Seq Kit v2 - Pico Input Mammalian, with rRNA depletion. […] sequenced on Illumina HiSeq 4000. […] (2) **V4**: […] Takara Bio SMART-Seq: SMART-Seq v4 Ultra Low Input RNA Kit, with PolyA selection for ribo depletion. […] sequenced on Illumina HiSeq 4000. […] (3) **TruSeq**: […] Stranded mRNA Sample Preparation Kit, with PolyA selection for ribo depletion. […] sequenced on Illumina HiSeq 2500.* […] *The experiment with the libraries prepared with the TruSeq kit was performed by Lahens et al.[11]. The Pico libraries were reproduced to repeat the experiment and ensure reproducibility*”. More information is available in their publication’s Methods section.

We obtained their count data (comprising 16184 genes observed in each of 18 samples) from the NCBI Gene Expression Omnibus (GEO; https://www.ncbi.nlm.nih.gov/geo/) with accession number GSE124167, which entails one count matrix per library preparation method. Then, we applied CONSTANd to each matrix separately, which means we let each library preparation method delineate an IPS. The distinction between ILB- and UNT-treated samples is considered a factor of biological interest. The same factors were specified in the model used by DESeq2. Any other remaining factors are ignored, because they would be confounded with a library preparation method.

### Human data (Li et al.)

This data set is a subset from the study by Li et al.[8], where standardized, commercial RNAs were sent to multiple sites (we consider sites L, R, V, W) for RNA-seq library preparation using ribo-depletion and/or poly-adenylation. A HiSeq 2500 was used for the libraries (both ribo-depletion and poly-adenylation) from site W; all other Illumina libraries (only ribo-depletion) were sequenced on a HiSeq 2000. “*Universal Human Reference RNA (740000, Agilent Technologies) and Ambion FirstChoice Human Brain Reference RNA (AM6000, Life Technologies) were used as the primary input RNAs for this study. These samples were labeled as […] A and B, respectively. […] External RNA Control Consortium (ERCC) “spike-in” synthetic transcripts were added at manufacturer recommended amounts (4456739, Life Technologies) to A and B standards*”. Then, they produced samples C and D by mixing a portion of samples A and B in 3:1 (C) and 1:3 ratios (D), respectively, as depicted in Figure 6. More information is available in their publication’s Online Methods section.

**Figure 6:**
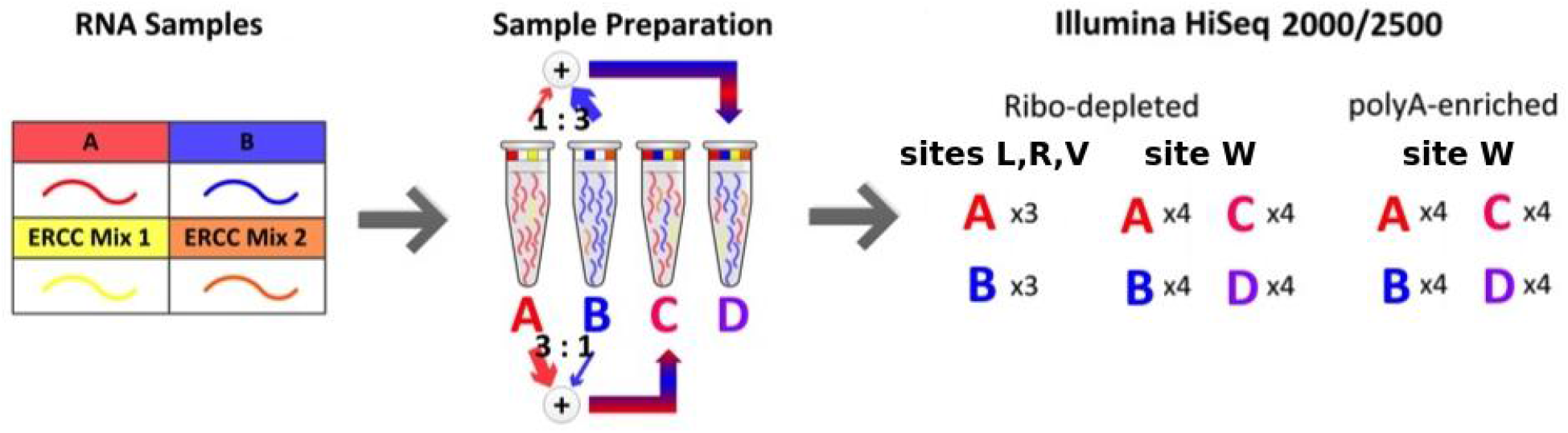
In the Human data set study, a total of 50 technical replicates of 4 types of samples (17xA, 17xB, 8xC, 8xD) were sequenced at 4 different sites using ribo-depletion-based libraries. At site W, some were also sequenced using a poly-adenylation-based library. Image source: adapted from Li et al.[8].

We obtained their count data (comprising 35779 genes observed in each of 50 samples) from GEO with accession number GSE48035, which entails one count table including all samples. We split that table into different tables according to sequencing site. For site W, we considered both the case where we left its site-specific table intact, as well as the case where we split it further into two tables according to library preparation method. As such, the IPSs are defined according to site and/or library preparation. We consider the distinction between the types of replicates A, B, C, D to be of biological interest. The same factors were specified in the model used by DESeq2. Any other remaining factors are ignored, because they would be confounded with either a site or a library preparation method.

## Abbreviations

IPS: identically processed (sub)set
IPFP: iterative proportional fitting procedure
GLM: generalized linear model

## Supplementary Information

### CONSTANd result without splitting Human data according to library preparation method

Figure S1 illustrates how a factor that introduces variability should delineate the IPSs. Even though all samples were collected with the same instrument in the same lab (W), the difference in library preparation protocol introduces enough variability to separate the samples according to library preparation in PC2. Hence, we should carefully take all such factors into account when defining our IPSs for normalization in a transcriptomics context in order to appropriately analyze the data.

**Figure S1:**
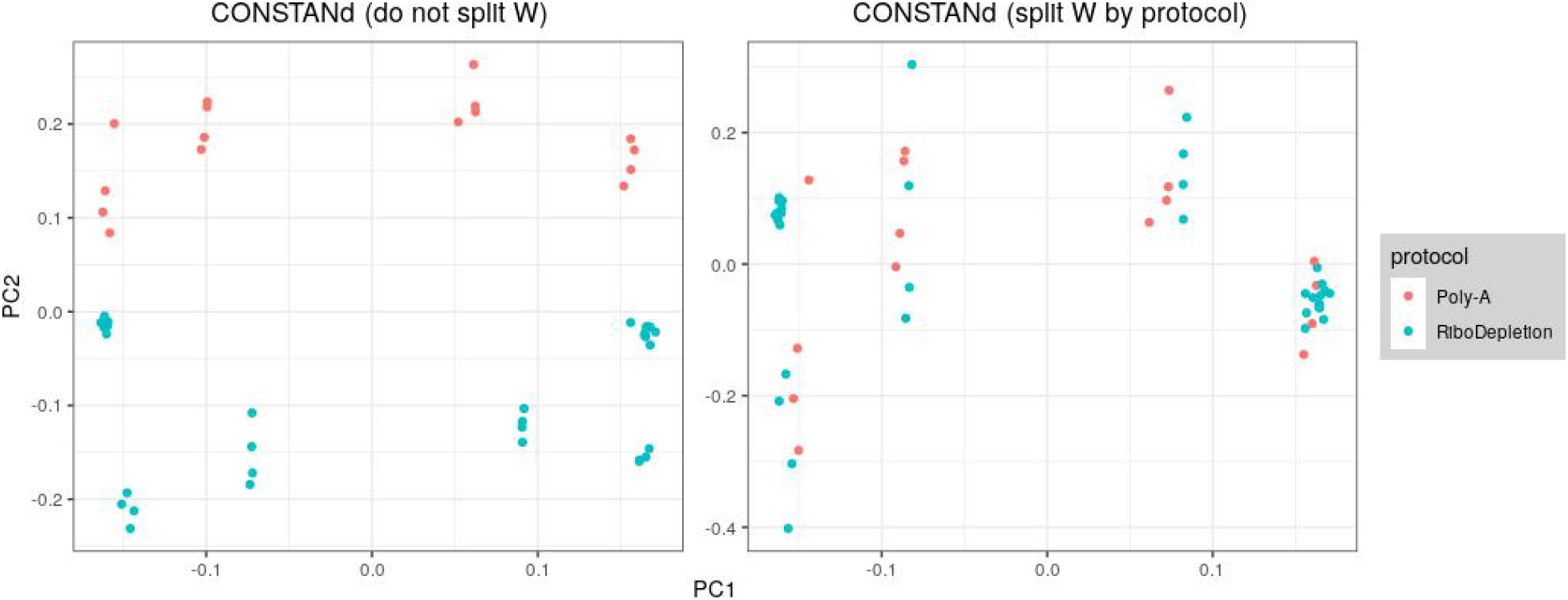
Splitting the count matrix of site W of the Human data set into two matrices based on the library preparation protocol and normalizing them separately will introduce a clear distinction between those samples in PC2.

### Sarantopoulou et al. did not use removeBatchEffects in DESeq2 analysis of Mouse data

Strangely enough, the clustering dendrogram from Sarantopoulou et al.[7] corresponds well to our dendrogram of the *original (un-normalized)* data. This is unexpected, since the following quote (clearly suggesting the data ought to be normalized) appears in their discussion of this figure: “*Hierarchical clustering by normalized expression correlation of all 18 samples shows clear distinction of the samples first by kit type and secondly by treatment. […] Thus the differences between the kits are more pronounced than the differences between the samples, in spite of a powerful treatment affecting thousands of genes with large effect sizes.*”

**Figure S2:**
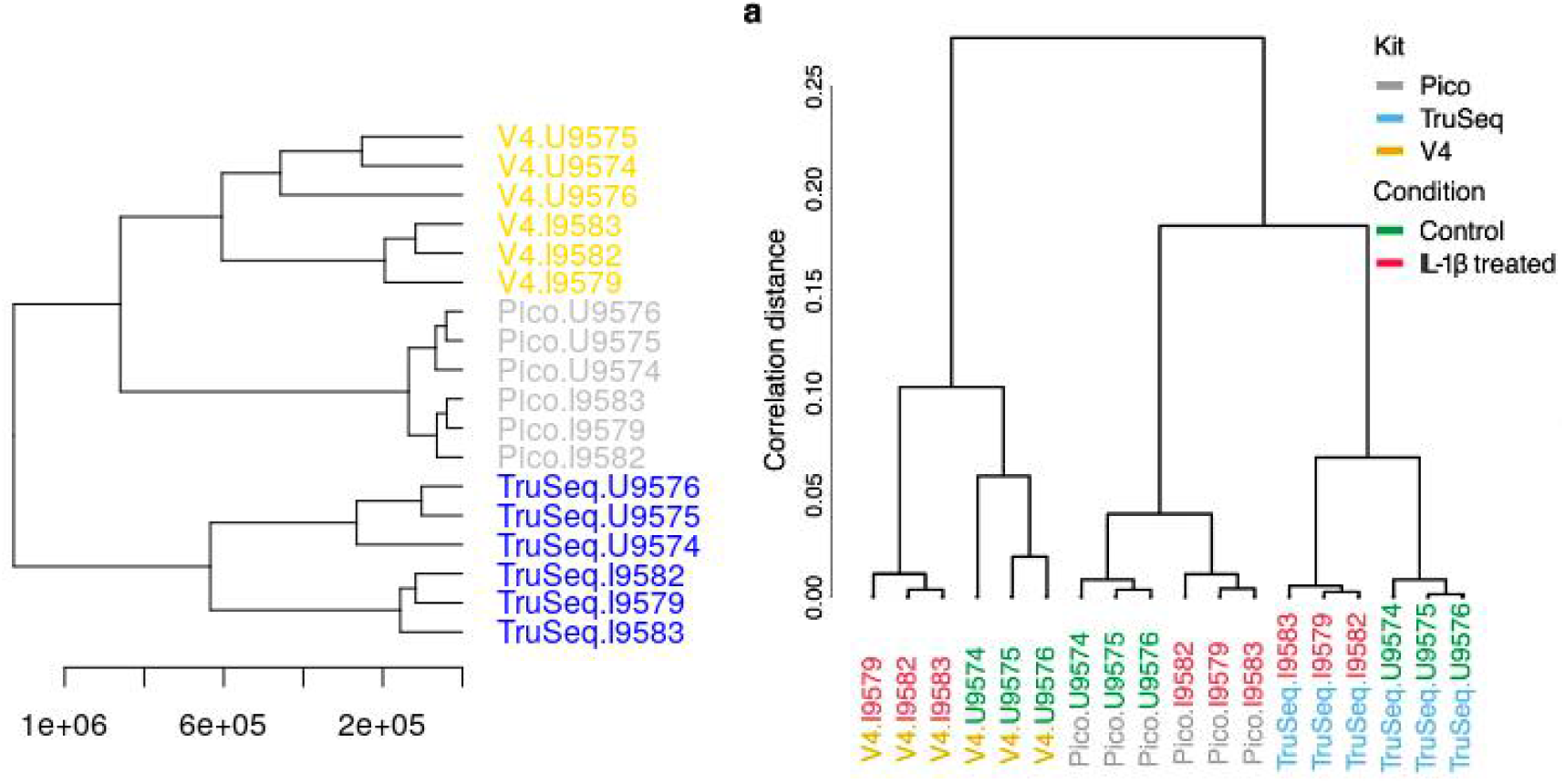
The hierarchical clustering (right) of the so-called normalized expression data taken directly from Sarantopoulou et al.[7] looks suspiciously similar to our clustering (left) of raw data.

Moreover, after our own DESeq2 (as well as CONSTANd) normalization we had obtained a very clear separation on PC1 due to biological condition, as shown in Fig. S3.

**Figure S3:**
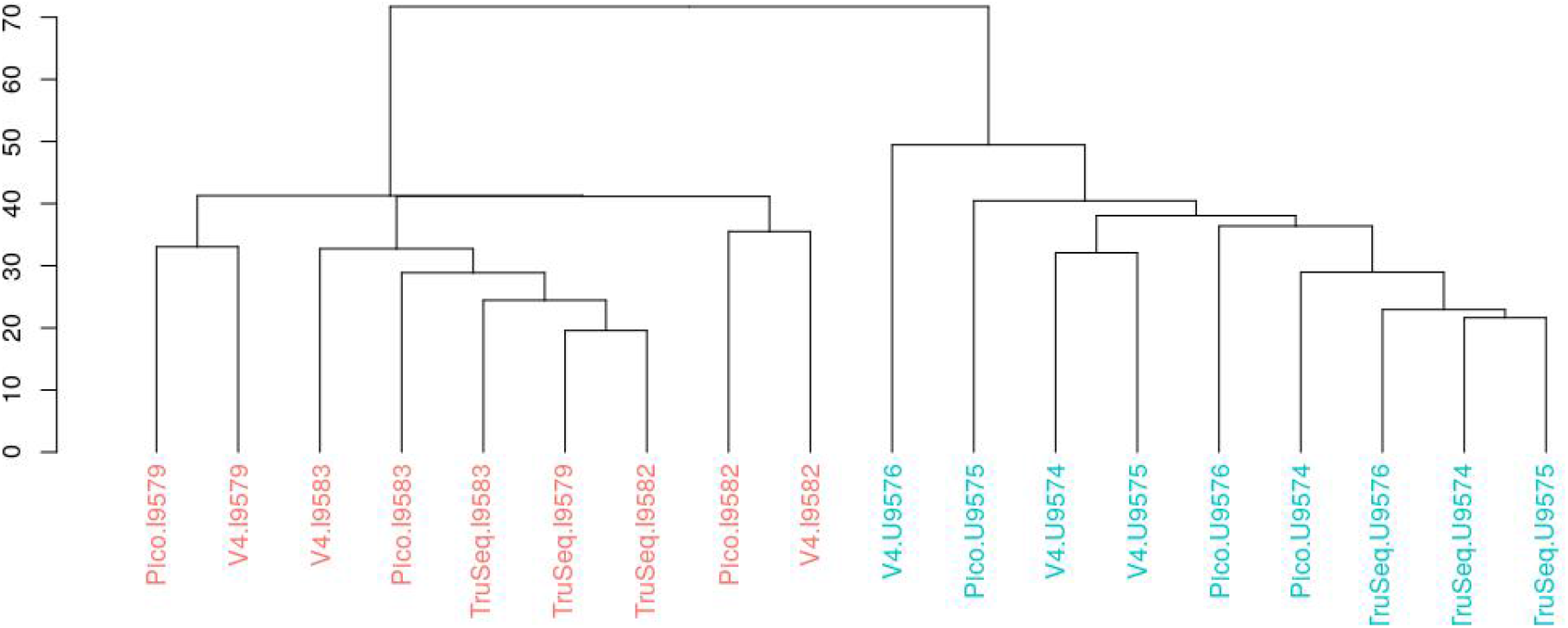
our hierarchical clustering of DESeq2-normalized data perfectly separates the samples based on biological treatment (ILB: red, UNT: blue).

This leads us to believe that the data depicted in the figure from Sarantopoulou’s paper has not had its kit effects removed by DESeq2’s removeBatchEffects function. We can double-check that last claim by reproducing Sarantopoulou’s result through *not* removing the kit effect ourselves. Indeed, in Fig. S4 we obtain a separation by kit on PC1. This putative mistake is a clear example suggesting that relatively simple and easy-to-use methods like CONSTANd may be preferred (at least in the triage phase) over more complicated and thus error-prone methods.

**Figure S4:**
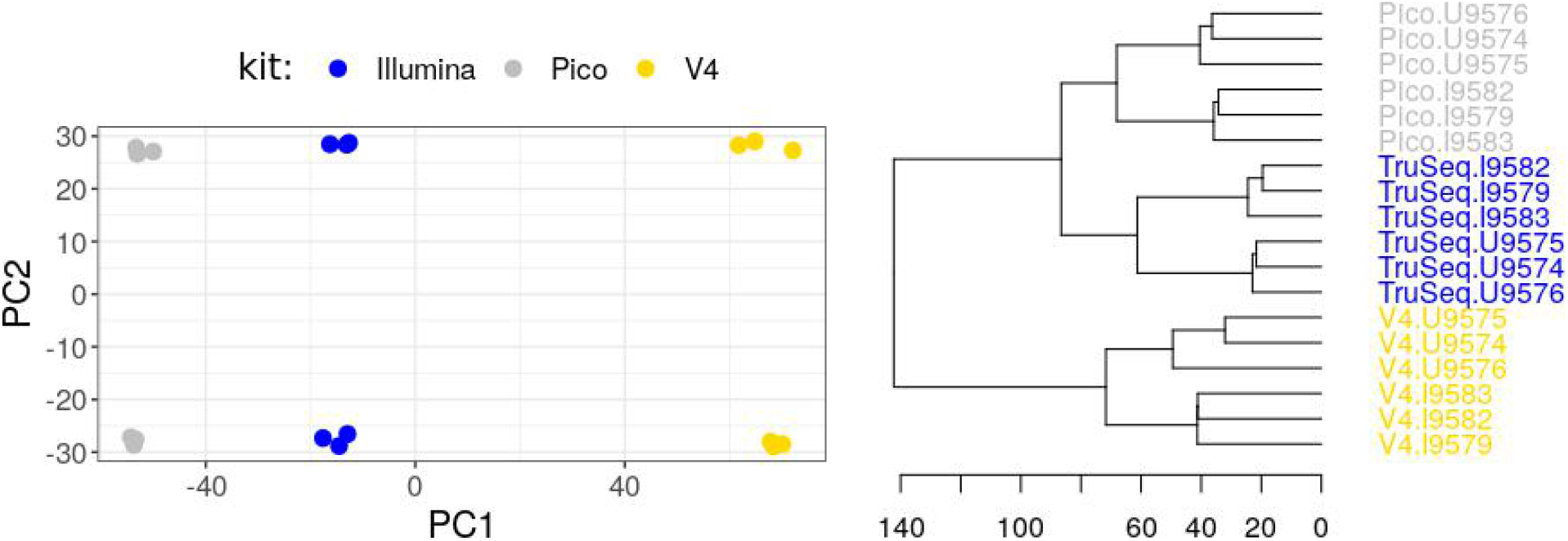
when not using removeBatchEffects after DESeq2, both the PCA plot and hierarchical clustering show the library preparation method as the primary source of variability.

### Additional figures

**Figure S5:**
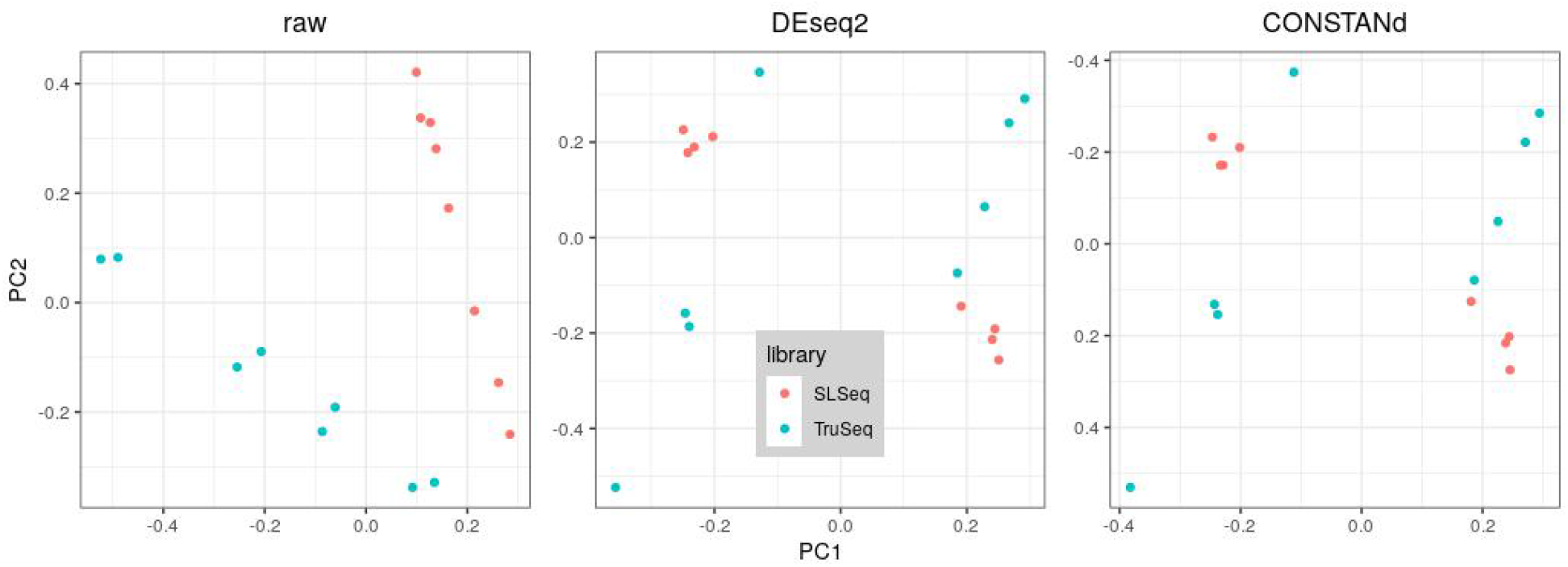
The PCA partially shows the raw data from the Leishmania data set is separable based on library preparation method. Both DESeq2 or CONSTANd normalization are able to largely remove this systematic effect.

**Figure S6:**
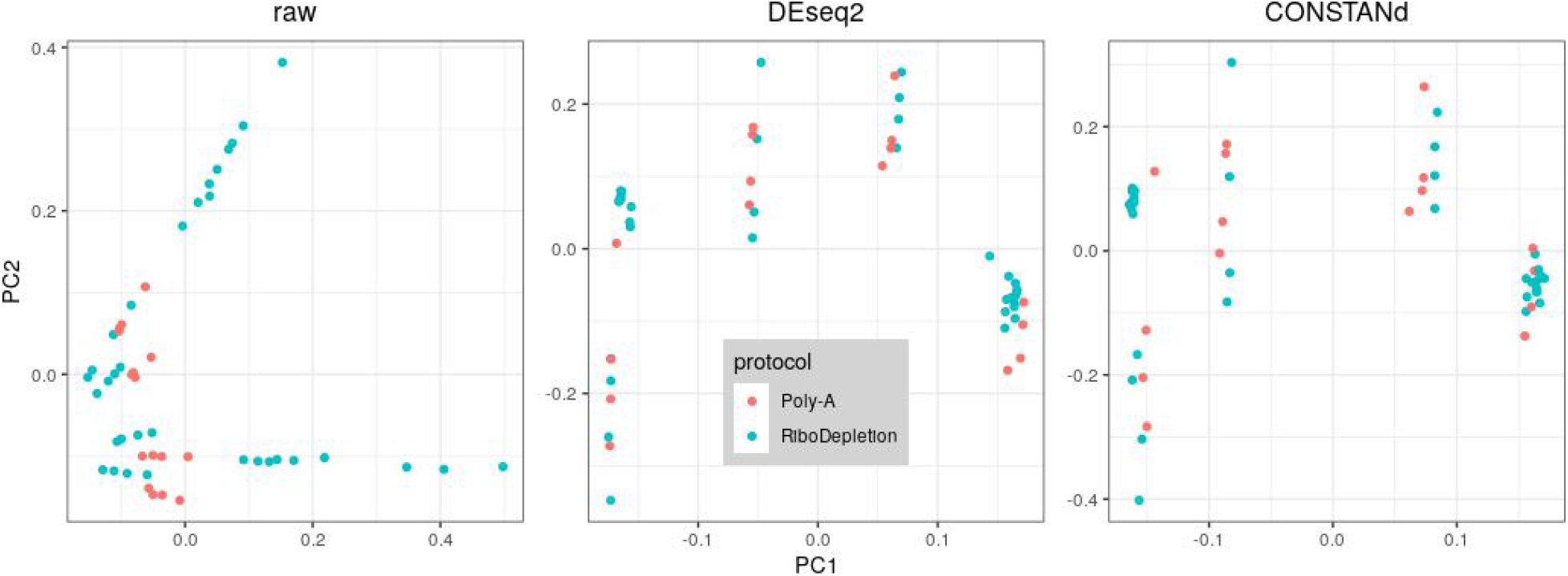
The PCA partially separates raw data from the Human data set based on library preparation protocol. Both DESeq2 or CONSTANd normalization are able to largely remove this systematic effect.

**Figure S7:**
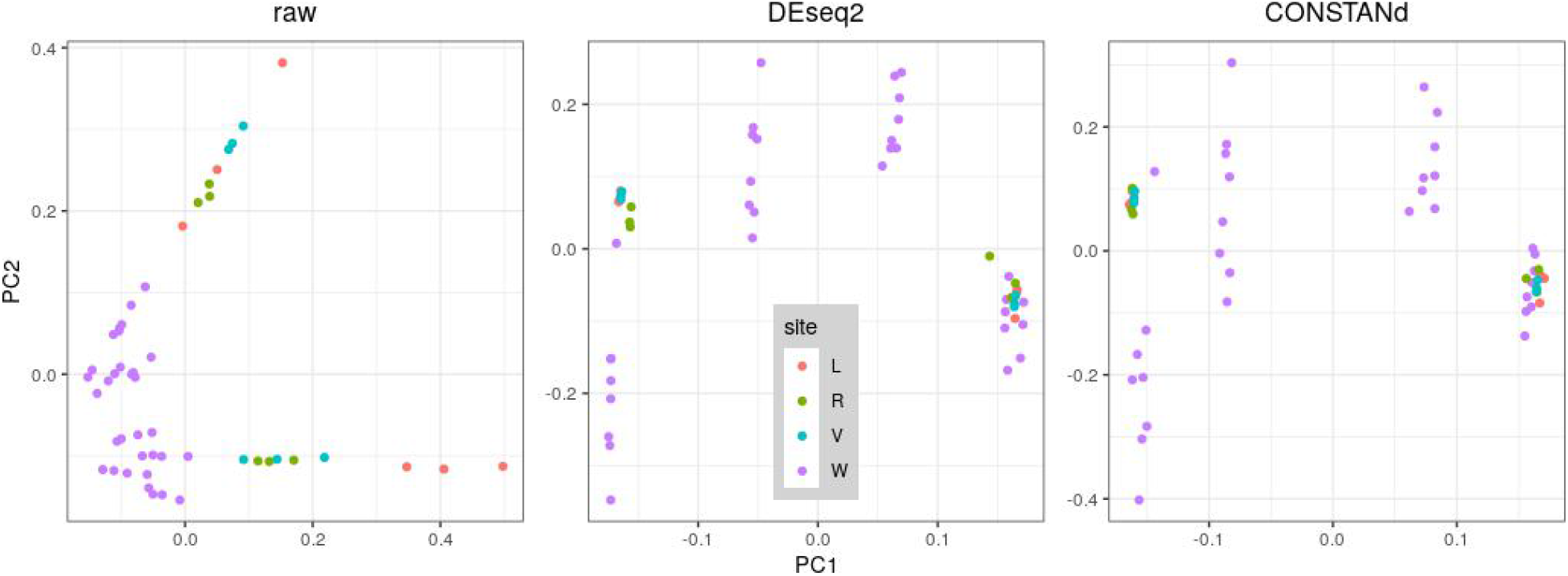
The PCA partially separates raw data from the Human data set based on site. Both DESeq2 or CONSTANd normalization are able to largely remove this systematic effect, except for the pooled tumor (A) samples from site W (purple, bottom-left; see Fig.5 in the main text).

**Figure S8:**
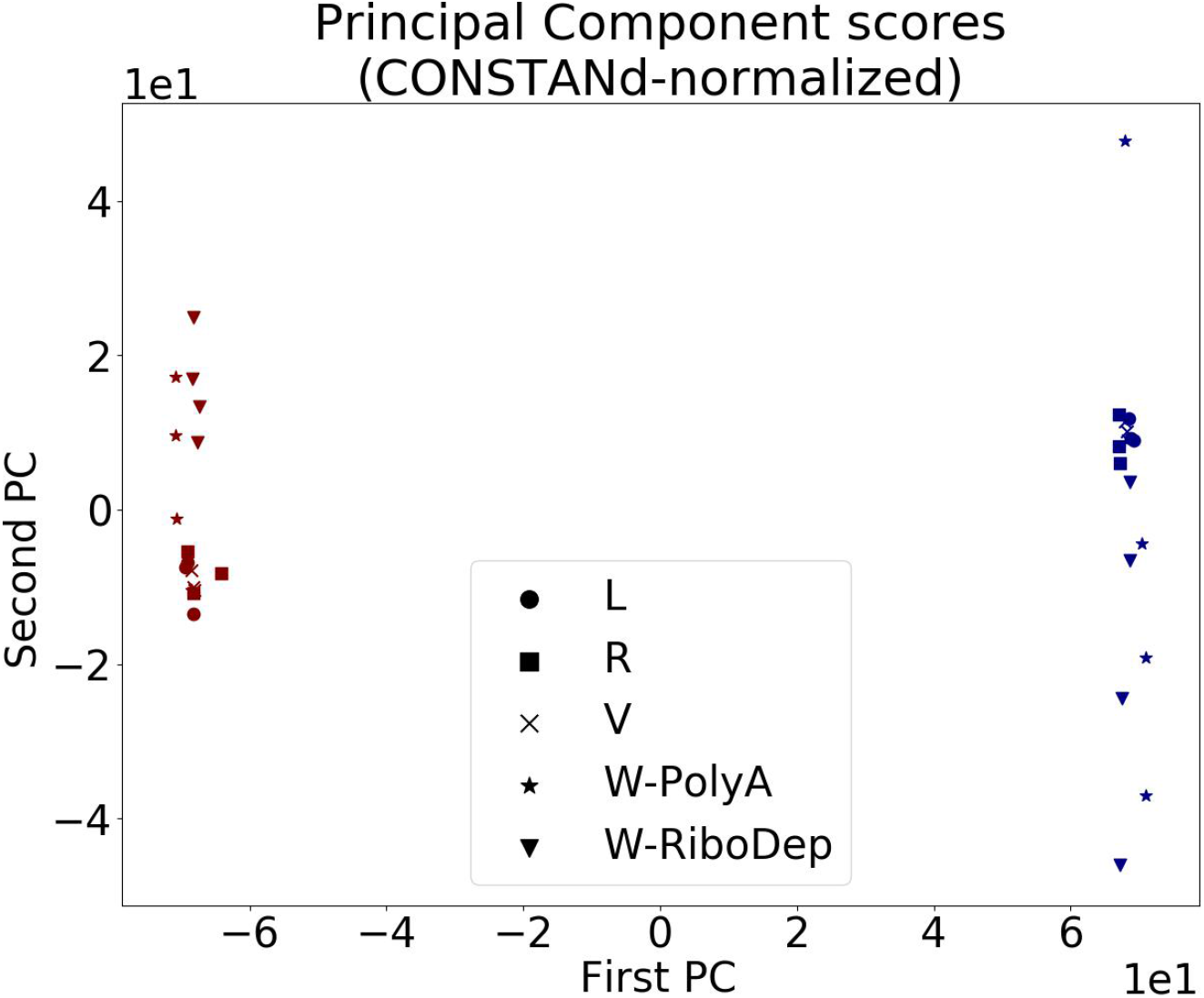
Even when considering only sample types A and B (omitting C and D to avoid confounding), the variability between samples from site W is still noticeably larger than those from other sites.

